# Mutations in *fbiD (Rv2983)* as a novel determinant of resistance to pretomanid and delamanid in *Mycobacterium tuberculosis*

**DOI:** 10.1101/2020.09.10.292508

**Authors:** Dalin Rifat, Si-Yang Li, Thomas Ioerger, Keshav Shah, Jean-Philippe Lanoix, Jin Lee, Ghader Bashiri, James Sacchettini, Eric Nuermberger

## Abstract

The nitroimidazole pro-drugs delamanid and pretomanid comprise one of only two new antimicrobial classes approved to treat tuberculosis (TB) in 50 years. Prior *in vitro* studies suggest a relatively low barrier to nitroimidazole resistance in *Mycobacterium tuberculosis*, but clinical evidence is limited to date. We selected pretomanid-resistant *M. tuberculosis* mutants in two mouse models of TB using a range of pretomanid doses. The frequency of spontaneous resistance was approximately 10^−5^ CFU. Whole genome sequencing of 161 resistant isolates from 47 mice revealed 99 unique mutations, 91% of which occurred in 1 of 5 genes previously associated with nitroimidazole activation and resistance: *fbiC* (56%), *fbiA* (15%), *ddn* (12%), *fgd* (4%) and *fbiB* (4%). Nearly all mutations were unique to a single mouse and not previously identified. The remaining 9% of resistant mutants harbored mutations in *Rv2983*, a gene not previously associated with nitroimidazole resistance but recently shown to be a guanylyltransferase necessary for cofactor F_420_ synthesis. Most mutants exhibited high-level resistance to pretomanid and delamanid, although *Rv2983* and *fbiB* mutants exhibited high-level pretomanid resistance, but relatively small changes in delamanid susceptibility. Complementing an *Rv2983* mutant with wild-type *Rv2983* restored susceptibility to pretomanid and delamanid. By quantifying intracellular F_420_ and its precursor Fo in overexpressing and loss-of-function mutants, we provide further evidence that Rv2983 is necessary for F_420_ biosynthesis. Finally, *Rv2983* mutants and other F_420_H_2_-deficient mutants displayed hypersusceptibility to some antibiotics and to concentrations of malachite green found in solid media used to isolate and propagate mycobacteria from clinical samples.

## INTRODUCTION

*Mycobacterium tuberculosis* remains the leading killer among infectious agents plaguing mankind, causing an estimated 1.45 million deaths in 2018 (1). The emergence and spread of rifampin-resistant (RR), multidrug-resistant (MDR) and extensively drug-resistant (XDR) *M. tuberculosis* makes tuberculosis (TB) control much more difficult. Detection of, and discrimination between, these forms of resistant TB requires laboratory confirmation of TB by rapid molecular test or culture and additional genotypic or phenotypic DST. Only approximately one-third of the estimated number of RR/MDR/XDR-TB cases are detected and initiated on treatment. Depending on the drug resistance profile, treatment has required administration of more toxic and less effective second- and third-line drugs for at least 9 months and up to 2 years (1, 2).

Delamanid and pretomanid are promising new bicyclic 4-nitroimidazole drugs that represent one of only two novel antimicrobial classes approved for clinical use against TB in 50 years. They have shown potential in pre-clinical and clinical studies to shorten and simplify the treatment of TB, including drug-resistant forms (3-10). Delamanid received conditional approval by the European Medicines Agency to treat MDR-TB in 2014 (11) but has had relatively limited clinical use to date. Pretomanid was recently approved by the U.S. Food and Drug Administration for treatment of MDR/XDR-TB as part of a novel oral, short-course regimen with bedaquiline and linezolid that produced favorable treatment outcomes in 90% of trial participants (4, 9).

Pretomanid and delamanid are prodrugs that require bioreductive activation of their aromatic nitro group by the mycobacterial 8-hydroxy-5-deazaflavin (coenzyme F_420_)-dependent nitroreductase Ddn in order to exert bactericidal activity (12). The reaction involves the transfer of two-electron hydride from the reduced form of cofactor F_420_ (F_420_H_2_) produced by an F_420_-dependent glucose-6-phosphate dehydrogenase (Fgd), the only enzyme known in mycobacteria to reduce F_420_ (13-15). Therefore, F_420_ biosynthesis and reduction by Fgd are essential for activation of delamanid and pretomanid. Three genes are known to be essential for F_420_ biosynthesis in *M. tuberculosis* complex (16, 17). *fbiC* encodes a 7,8-didemethyl-8-hydroxy-5-deazariboflavin (Fo) synthase that catalyzes the condensation of 5-amino-6-ribitylamino-2,4 (1*H*, 3*H*)-pyrimidinedione and tyrosine to form the F_420_ precursor Fo (18, 19). *fbiA* encodes a transferase that is now known to catalyze the transfer of a phosphoenolpyruvyl moiety to Fo to generate dehydro-F_420_-0, while *fbiB* encodes a bifunctional enzyme that reduces dehydro-F_420_-0 and then catalyzes the sequential addition of a variable number of glutamate residues to F_420_-0 to yield coenzyme F_420_-5 or -6 in mycobacteria (20). A fourth gene, MSMEG_2392, was shown to be necessary for F_420_ synthesis, but not Fo synthesis, in *Mycobacterium smegmatis* (21). Its homologue in *M. tuberculosis, Rv2983*, was recently cloned and purified to perform an *in vitro* assay to generate dehydro-F_420_-0 by using purified Rv2983 and FbiA, GTP, phosphoenolpyruvate (PEP) and Fo followed by binding study of PEP with the crystallized Rv2983, which proved Rv2983 to be aPEP guanylyltransferase (designated FbiD) that synthesizes the phosphoenolpyruvyl moiety that is subsequently transferred to Fo by FbiA (20). However, whether *Rv2983*, now known as *fbiD*, is essential for F_420_ biosynthesis in *M. tuberculosis* awaits confirmation. Furthermore, it remains unknown whether *Rv2983* is necessary for the activation of pretomanid and delamanid.

A variety of loss-of-function mutations in *ddn, fgd* and *fbiA-C* causing delamanid and pretomanid resistance are readily selected *in vitro* in *M. tuberculosis* complex (17, 19, 22-24). However, a recent study found that 17% of the resistant isolates selected *in vitro* by pretomanid harbored no mutations in these genes (23). Furthermore, there has been no comprehensive study of evolution of resistance *in vivo* during treatment with either agent. Given that clinical usage of delamanid and pretomanid is increasing and fitness costs arising from resistance mutations may differ between *in vitro* and *in vivo* conditions, the paucity of data relating to the emergence of resistance *in vivo* is alarming. Therefore, we set out to study bacterial genetic, host and pharmacological factors associated with emergence of nitroimidazole resistance in two murine models of TB. In so doing, we identified loss-of-function mutations in *Rv2983* as a novel determinant of pretomanid and delamanid cross-resistance and proved its essentiality for F_420_ biosynthesis in *M. tuberculosis*, findings that support its role as FbiD in the recently revised F_420_ biosynthesis pathway. We also characterized additional phenotypes of the *Rv2983* mutants, showing them to be hypersensitive to some stress conditions and antibiotics, and to malachite green (MG), an organic compound used as a selective decontaminant in solid media for culturing *M. tuberculosis*. The latter finding raises important concerns that isolation and propagation of nitroimidazole-resistant mutants from clinical samples may be adversely affected by use of some MG-containing media, such as Lowenstein-Jensen media. Together, these findings have important implications for the development of both genotypic and phenotypic methods for detection of nitroimidazole resistance in clinical samples.

## RESULTS

### Spontaneous pretomanid-resistant mutants exist at a relatively high frequency in infected mice and are selectively amplified by treatment with active doses of pretomanid

To study the dose-response of pretomanid and explore the genetic spectrum of nitroimidazole resistance selected *in vivo*, we established chronic *M. tuberculosis* infections in mice and then treated with a range of pretomanid doses spanning the clinical exposure range for up to 8 weeks. Because the lungs of TB patients feature a heterogeneous array of lesion types resulting in diverse microenvironments and pharmacological compartments that alter the drug susceptibility and drug exposure of resident tubercle bacilli (25, 26), we used both C3HeB/FeJ mice, which develop caseating lung lesions in response to infection, and BALB/c mice, which do not, to investigate the impact of host pathology on mutant selection. Despite lower CFU counts on the day after infection (W-8) in C3HeB/FeJ mice (1.67 log_10_ CFU per lung) compared to BALB/c (2.26 log_10_) (*p* <0.001), higher CFU counts were observed in C3HeB/FeJ mice 8 weeks later, on the day treatment started (D0), and after 3 weeks of treatment in almost all groups (*p* <0.001 - 0.05) (Fig. 1A), consistent with the greater susceptibility of this strain to *M. tuberculosis* infection. Three C3HeB/FeJ mice treated with 1000 mg/kg required euthanasia during the second week of treatment, prompting a dose reduction from 1000 mg/kg to 600 mg/kg in both strains. Nevertheless, a clear pretomanid dose-response relationship was observed in both mouse strains after 3 weeks of treatment (Fig. 1A). The three remaining C3HeB/FeJ mice treated with 600 mg/kg beyond the week 3 time point were euthanized after 5 weeks of treatment due to toxicity. One had no detectable CFU and two had ≤ 2.0 log_10_ CFU of pretomanid-resistant *M. tuberculosis*. After 8 weeks of treatment, total CFU counts fell in a dose-dependent manner in BALB/c mice before a plateau was reached at doses ≥ 300 mg/kg, where resistant CFU were higher and replaced the susceptible CFU (*p* < 0.05) (Fig. 1B). Spontaneous pretomanid-resistant CFU comprised approximately 10^−5^ of the total CFU in the absence of drug pressure in untreated BALB/c mice and the proportion of the total CFU that was comprised of pretomanid-resistant CFU increased with dose up to the 300 mg/kg dose group. Dose-dependent bactericidal activity was also observed in C3HeB/FeJ mice (Fig. 1C). However, selective amplification of pretomanid-resistant mutants was more extensive and occurred at lower doses than in BALB/c mice (Fig. 1B and 1C). We were not able to measure the spontaneous frequency of resistant mutants in untreated C3HeB/FeJ mice because they succumbed to infection prior to week 8. Pretomanid-resistant CFU replaced susceptible CFU in C3HeB/FeJ mice receiving doses as low as 30 mg/kg and pretomanid-resistant CFU counts were roughly 10 times higher in C3HeB/FeJ mice compared to BALB/c mice (Fig. 1B and C), which indicates greater potential for selective amplification of pretomanid resistance with monotherapy in this strain. Most resistant isolates grew on plates containing 10 µg/ml of pretomanid, but some had fewer CFU on plates containing 10 µg/ml than on those containing 1 µg/ml of pretomanid.

**Fig. 1.**
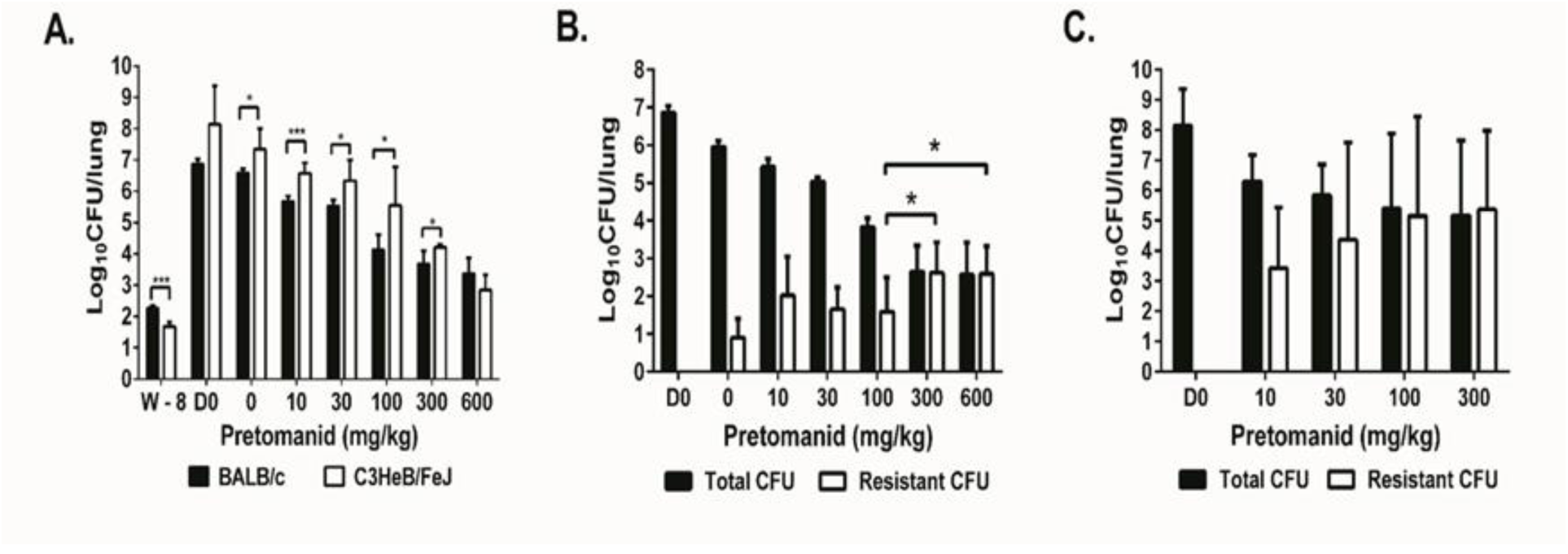
Selective amplification of spontaneous pretomanid-resistant mutants during pretomanid monotherapy in mice is dose-dependent and is more pronounced in C3HeB/FeJ mice. After aerosol infection with *M. tuberculosis* H37Rv, BALB/c and C3HeB/FeJ mice were treated with a range of doses of pretomanid for 8 weeks and sacrificed at different time points before and after treatment for lung CFU counts. A. Mean (± S.D.) total lung CFU counts on the day after infection (W-8), on the day of treatment initiation (D0), and after 3 weeks of treatment with the indicated pretomanid dose (in mg/kg body weight). Dose-dependent bactericidal activity was observed in both strains; B. Mean (± S.D.) total and PMD-resistant lung CFU counts in BALB/c mice on day 0 and after 8 weeks of treatment with the indicated pretomanid dose. Dose-dependent bactericidal activity and selection of PMD-resistant bacteria was observed, with the resistant population overtaking the susceptible population at doses ≥ 300 mg/kg; C. Mean (± S.D.) total and PMD-resistant lung CFU counts in C3HeB/FeJ mice on day 0 and after 8 weeks of treatment with the indicated pretomanid dose. Dose-dependent bactericidal activity and selection of PMD-resistant bacteria was observed, with the resistant population overtaking the susceptible population at doses ≥ 30 mg/kg. *\* p <* 0.05, *\*\*\* p <* 0.001

### Whole genome sequencing of pretomanid-resistant mutants revealed diverse mutations in *Rv2983* or in one of five other genes known to be required for pretomanid activation

To characterize mutations associated with pretomanid resistance *in vivo*, we performed WGS on 136 individual pretomanid-resistant colonies and 25 colony pools picked from 47 individual mice harboring pretomanid-resistant CFU after 8 weeks of treatment (Table S2-S4). Each individual isolate had an isolated mutation in one of the 5 genes previously shown to be required for pretomanid activation or in *Rv2983*, a gene not previously associated with nitroimidazole resistance. Overall, 99 unique and diverse mutations in these 6 genes were identified. Each mouse lung harbored 1 to 4 unique mutations. Except for a few mutations (K9N (*fgd*), R322L (*fbiC*) and Q120P (*fbiA*)) shared by two mice each, no two mice harbored the same mutation. Moreover, comparing the 99 unique mutations identified in our study with the 151 unique mutations in the 5 previously recognized genes selected *in vitro* (23), only 4 mutations were found in the same position and only the W79 stop (*fbiA*) and N336K (*fbiC*) mutations were found in both datasets. In our pooled samples, mutations in all the genes mentioned above were detected except mutations in *fbiB* (Table S5). Taken together, these data reveal a remarkably large “target size” for chromosomal mutations conferring resistance to nitroimidazole pro-drugs *in vivo* as well as *in vitro*.

In both BALB/c and C3HeB/FeJ mice, more than half of the resistant isolates were *fbiC* mutants (56%) (Fig. 2A). For the other five genes, the rank order by mutation frequency was *Rv2983* (13%) and *fbiA* (13%) > *ddn* (9%) > *fbiB* (6%) > *fgd* (4%) in BALB/c mice (Fig.2B and Table S5) and *fbiA* (18%) > *ddn* (16%) > *fgd* or *Rv2983* (4%) > *fbiB* (2%) in C3HeB/FeJ mice (Fig.2C and Table S5). No significant differences in mutation frequencies between BALB/c and C3HeB/FeJ mice were observed, although a trend towards more *Rv2983* mutations in BALB/c mice (7/54, 13% of all mutations) compared to C3HeB/FeJ mice (2/45, 4%) was detected. The mutations identified in *Rv2983* included 8 point mutations resulting in the following amino acid substitutions: R25S, R25G, A68E, A132V, G147C, C152R, Q114R and A198P, as well as an insertion of C after A27 and a deletion of I129 (-ATC) (Tables S2-S4). There were no clear associations between pretomanid dose or concentration and the mutated gene. Mutations in *fbiC* comprised a higher proportion of those selected in our *in vivo* study compared to the proportion selected in a previous in *vitro* study (26%, *p* = 0.0001) (23), implying that such mutants may have superior fitness *in vivo* relative to other mutants. On the other hand, mutations in *ddn* (29%) were more frequent after *in vitro* selection than in our mouse models (12%) (*p* =0.001). *In vitro* mutation frequencies for *fbiA, fgd* and *fbiB* (19%, 7% and 2%, respectively) were similar to our findings in mice.

**Fig. 2.**
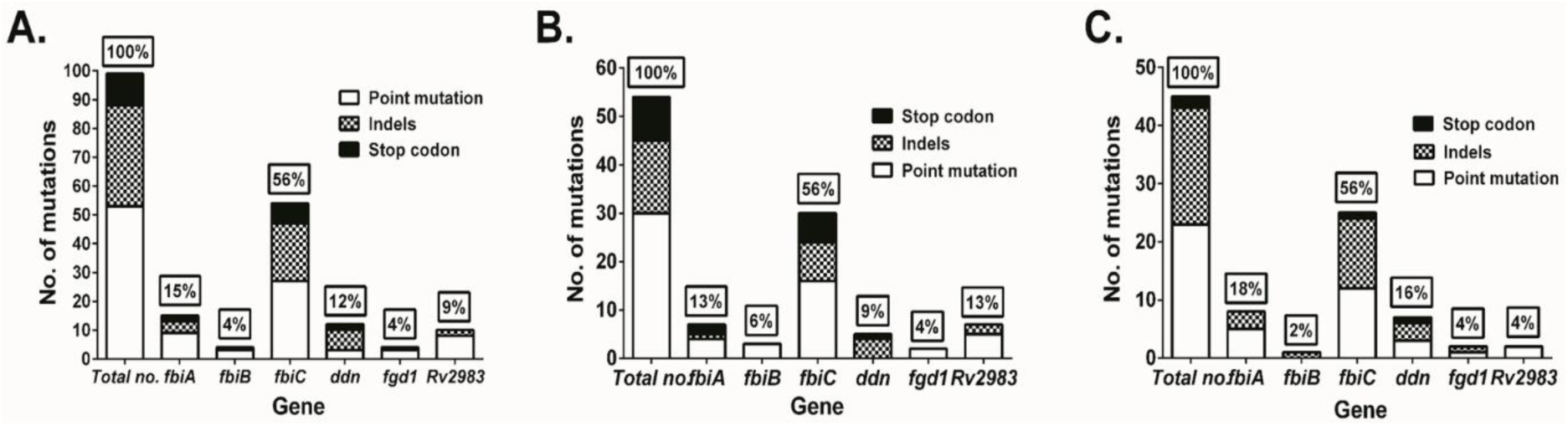
Mutation frequencies and mutation types of genes associated with pretomanid resistance. WGS was performed with 136 pretomanid-resistant colonies and 25 colony pools picked from 47 individual mice harboring pretomanid-resistant CFU after 8 weeks of treatment. 99 unique mutations in these 6 genes were identified. A. Overall mutation frequencies; B. Mutation frequencies and mutation types in BALB/c mice; C. Mutation frequencies and mutation types in C3HeB/FeJ.

Among the 99 unique mutations, all but one (an IS6110 insertion located in 85-bp upstream of the *fbiC* coding sequence in isolate KA-026a) (Table S3) were found within the coding regions of the six genes. In total, 54% (53/99) were non-synonymous point mutations (no synonymous point mutations were identified), 35% (35/99) were insertions or deletions (indels), and 11% (11/99) were substitutions resulting in a new stop codon (Fig.2A and Table S5). No significant difference in the distribution of point mutations and indels was found between ours and the *in vitro* study by Haver, *et al*. However, the frequency of stop codon substitutions in the latter study (26%, 40/151) was higher than that observed in the present study (11%, 11/99) (*p* = 0.004), 85% (34/40) of which were in *ddn* in the latter study (23).

### Mutations in *Rv2983* cause resistance to pretomanid and delamanid

To prove that loss-of-function mutations in *Rv2983* are sufficient for nitroimidazole resistance, merodiploid complemented strains were constructed by introducing a copy of the wild type *Rv2983* gene into B101, an *Rv2983* mutant (A198P), through site-specific integration (27, 28). Susceptibility testing confirmed significantly higher nitroimidazole MICs against the *Rv2983* mutant and full restoration of susceptibility in the complemented strains. Remarkably, however, the upward shift in pretomanid MIC (i.e., >128x) associated with this mutation was significantly greater than the shift in delamanid MIC (i.e., 8x), and the delamanid MIC of 0.06 µg/ml against the *Rv2983* mutant was the same as the recommended critical concentration for susceptibility testing in MGIT medium (29). MICs were determined against additional isolates with mutations in each of the 6 genes (Table 1). Interestingly, whereas mutations in *fbiC, fbiA, fgd* and *ddn* were often associated with high-level resistance to both pretomanid and delamanid, both *Rv2983* and *fbiB* mutants exhibited high-level pretomanid resistance while delamanid MICs hovered around the MGIT breakpoint of 0.06 µg/ml, or more than 100 times lower than delamanid MICs against *ddn* mutants and most other *fgd* and F_420_ biosynthesis mutants. The sole exception was an *Rv2983* frameshift mutant (BA019a) that exhibited high-level resistance to both compounds.

**Table 1.** Pretomanid and delamanid MICs against the parent H37Rv strain and isogenic mutants selected in mice.

| Name of isolate selected | Mutated gene | Mutation | Pretomanid MIC (µg/ml) | Delamanid MIC (µg/ml) |
| --- | --- | --- | --- | --- |
| H37Rv | n/a | n/a | 0.25 | 0.008 |
| BA019a | *Rv2983* | +C in aa 27 | >32 | >16 |
| BA032 | *Rv2983* | G147C | >32 | 0.06 |
| BA043 | *Rv2983* | A132V | >32 | 0.06 |
| BA060 | *Rv2983* | -ATC in aa 129 | >32 | 0.06 |
| BA078-82 | *Rv2983* | R25S | >32 | 0.03-0.06 |
| B101 | *Rv2983* | A198P | >32 | 0.06 |
| BA120 | *Rv2983* | C152R | >32 | 0.06 |
| KA003 | *Rv2983* | A68E | >32 | <0.03 |
| KA016 | *Rv2983* | Q114R | 16- >32 | 0.06-0.125 |
| BA026a | *fbiB* | L15P | 8-32 | 0.06-0.125 |
| BA069 | *fbiB* | W397R | 16 | 0.03 |
| BA070 | *fbiB* | L173P | 32 | 0.125 |
| KA006 | *fbiB* | -T in aa 684 | 32 | 0.06-0.125 |
| BA074 | *fbiA* | S219G | 32 | 0.03 |
| BA084 | *fbiA* | Q27* | >32 | >16 |
| KA043 | *fbiA* | D49G | >32 | >16 |
| KA058 | *fbiA* | -G in aa 47 | >32 | >16 |
| KA067 | *fbiA* | L308P | >32 | >16 |
| KA085 | *fbiA* | Q120P | 32 | >16 |
| KA096 | *fbiA* | D286A | >32 | >16 |
| BA017 | *fbiC* | C562W | >32 | >16 |
| BA035 | *fbiC* | R25G | 16-32 | 0.03 |
| BA075 | *fbiC* | M776T | 16-32 | 0.03 |
| KA004 | *fbiC* | G194D | >32 | 1 |
| KA014 | *fbiC* | -C in aa 20 | >32 | 2 |
| KA017 | *fbiC* | K684T | >32 | >16 |
| KA026a | *fbiC* | IS6110 ins. 85bp upstream of *fbiC* | >32 | >16 |
| KA031 | *fbiC* | L377P | >32 | >16 |
| KA073 | *fbiC* | A827G | >32 | >16 |
| BA002 | *fgd* | K9N | 32 | 0.5 |
| KA050 | *fgd* | G191D | >32 | >16 |
| KA088 | *ddn* | R112W | ≥32 | >16 |
| KA091 | *ddn* | IS6110 ins. in D108 | ≥32 | >16 |
| KA093 | *ddn* | -G in aa 39 | >32 | >16 |

### *Rv2983* is required for F_420_ biosynthesis

To demonstrate that *Rv2983* is required for F_420_ biosynthesis, we measured the production of Fo and F_420_ in *M. smegmatis* strains overexpressing *Rv2983* and in *M. tuberculosis Rv2983* mutant strains compared with their corresponding control strains. Previous studies in *Mycobacterium bovis* BCG showed no detected Fo or F_420_ in an *fbiC* mutant, detected Fo with no detected F_420_ in an *fbiA* mutant or with only a small amount of F_420_-0 detected only in an *fbiB* mutant (17, 19). In the current study, *Rv2983* was cloned into the IPTG-inducible expression vector pYUBDuet and p*fbiC* (designated p*Rv2983* and p*fbiC*-*Rv2983*, respectively), followed by successful transformation of *M. smegmatis*, along with pYUBDuet and p*fbiABC* controls. Relative fluorescence was assessed in these strains compared to the control strain containing the empty vector pYUBDuet. Overexpression of *Rv2983 in M. smegmatis* increased F_420_ production but resulted in little change in Fo production compared to the control strain after 6 and 26 hours of induction with IPTG (Figs. 3A and 3B). As expected, mutation of *Rv2983* in the *M. tuberculosis* B101 mutant markedly reduced F_420_ production, resulting in accumulation of Fo relative to the wild-type.

**Fig. 3.**
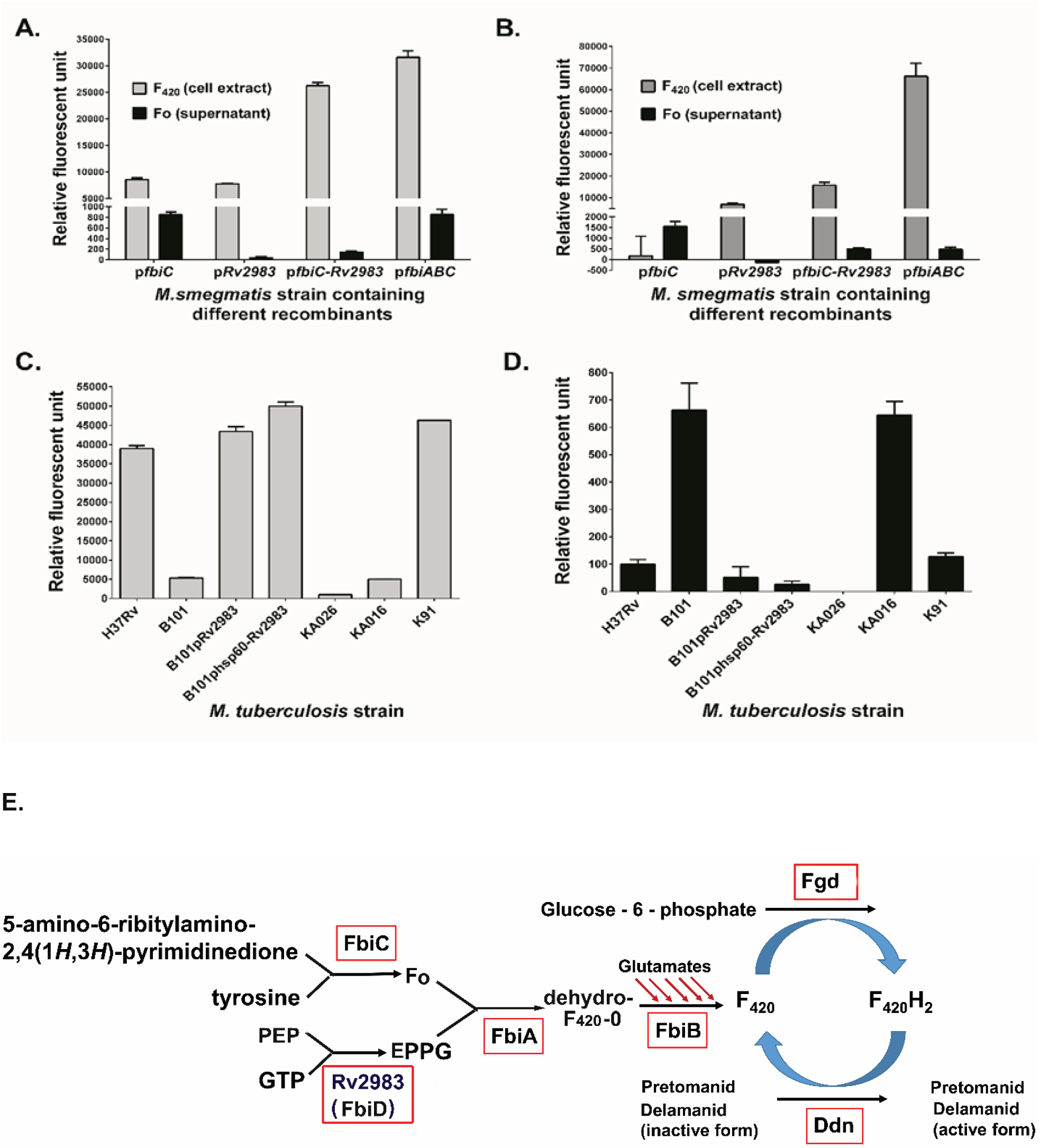
Rv2983 is required for efficient F_420_ synthesis from Fo. F_420_ and Fo content measured in *M. smegmatis* strains harboring different recombinants relative to the control strain containing the empty vector pYUBDuet after 6 (A) and 26 (B) hours of 1mM IPTG induction; F_420_ (C) and Fo (D) content was measured in the *Rv2983* mutant strains of *M. tuberculosis* and the control strains including B101 (Δ*Rv2983*, A198P), KA016 (Δ*Rv2983*, Q114R), H37Rv (wild-type), B101 complemented strain (pMH94-Rv2983), B101 complemented strain (pMH94-hsp60-*Rv2983*), KA026 (Δ*fbiC*, IS6110 insertion in 85-bp upstream of *fbiC*), and K91 (Δ*ddn*, IS6110 insertion in aa D108), after growth in 7H9 broth for 6 days. Schematic diagram (E) of proposed nitroimidazole activation pathway showing Rv2983 as FbiD catalyzing EPPG biosynthesis. Fo, 7,8-didemethyl-8-hydroxy-5-deazariboflavin; PEP, phosphoenolpyruvate; GTP, guanosine triphosphate; EPPG, enolpyruvyl-diphospho-5’-guanosine.

Complementation fully restored the wild-type phenotype (Figs. 3C and 3D). Overexpression of *fbiC* in *M. smegmatis* increased Fo and, consequently, F_420_ concentrations, as expected. Relative to the control strain, F_420_ concentrations were similar when either *fbiC* or *Rv2983* was over-expressed alone (Fig. 3A). Interestingly, when *Rv2983* was co-overexpressed with *fbiC*, a dramatic increase in F_420_ was observed relative to over-expression of either gene alone (3.4 and 3.1-fold, respectively) after 6 hours of IPTG induction (*p*<0.001), with corresponding significant decreases of Fo levels after 6 and 26 hours of IPTG induction (5.8 and 3.1-fold; *p*<0.005 and 0.05, respectively), which were similar to the results of co-overexpression of *fbiA, fbiB and fbiC* as a positive control (Fig. 3A and 3B). These results suggest that the excess Fo produced by *fbiC* over-expression was efficiently converted to F_420_ by over-expressed Rv2983. On the other hand, although a small amount of F_420_ was observed in cell extracts of two *Rv2983* point mutants (B101 [A198P] and KA016 [Q114R]), their F_420_ content was significantly lower than that of the wild type (7.3 and 7.7-fold) (*p* < 0.001) and complemented B101 mutant (Fig. 3C). As expected, Fo accumulated in the two *Rv2983* mutant strains relative to the wild-type (6.7 and 6.5-fold; *p*<0.05 and 0.005, respectively) (Fig. 3D), indicating that Fo was not efficiently converted to F_420_ in the presence of a mutated *Rv2983*. Two other pretomanid-resistant strains were also assessed as controls. The KA026 mutant with an IS6110 insertion 85 bp upstream of *fbiC* had undetectable Fo and very little F_420_ content, while the KA91 mutant with an IS6110 insertion at amino acid position 108 of Ddn showed a wild-type phenotype with respect to F_420_ and Fo concentrations (Figs. 3C and D).

### *Rv2983* is necessary for resistance to oxidative stress and progressive hypoxia but not for growth and survival in BALB/c mice

An F_420_-deficient *fbiC* mutant of *M. tuberculosis* was previously shown to be hypersensitive to oxidative stress (30, 31). To investigate the importance of Rv2983 under oxidative stress, the wild-type H37Rv, the *Rv2983* mutant (B101) and two complemented strains (B101-p*Rv2983* and B101-p*hsp60-Rv2983*) were exposed to 20 µM and 100 µM of menadione. In the absence of menadione, no significant difference in growth kinetics was observed between strains (Fig. S4), confirming that *Rv2983* and F_420_ are dispensable for growth in nutrient-rich 7H9 broth. However, the *Rv2983* mutant was markedly more susceptible to menadione (Fig. 4A-B).

**Figure 4.**
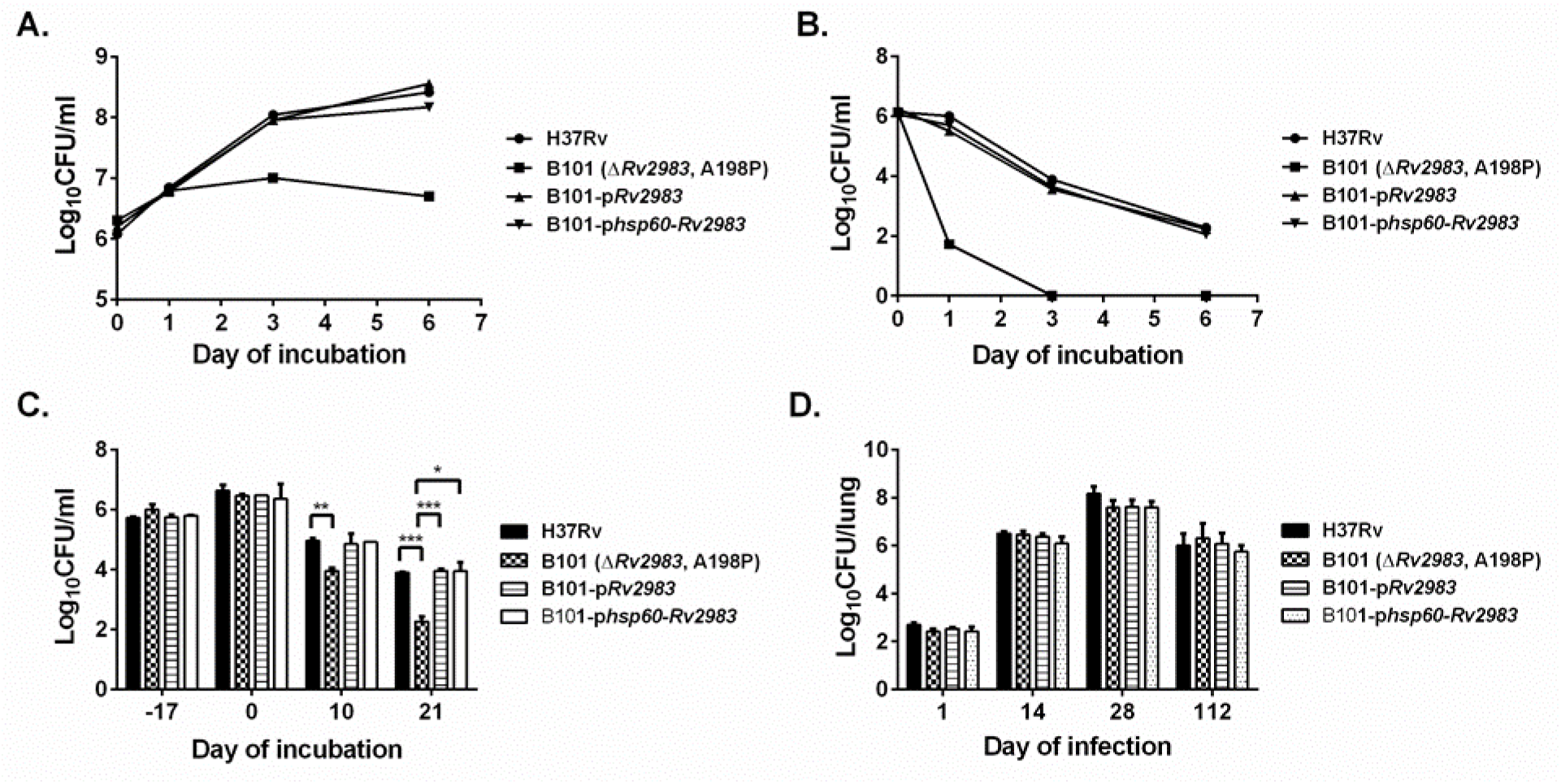
F_420_-deficient pretomanid-resistant *Rv2983* mutant is hypersensitive to oxidative stress and progressive hypoxia, but is not attenuated in BALB/c mouse lungs. A. *Mtb* growth kinetics in 7H9 broth containing 20 µM menadione; B. *Mtb* growth kinetics in 7H9 broth containing 100 µM menadione; C. *Mtb* growth and survival under progressive hypoxia; D. Lung CFU counts in BALB/c mice after aerosol infection with *Mtb* strains.

*M. tuberculosis* encounters hypoxia and enters a state of non-replicating persistence in closed caseous foci in diseased lungs (32). In order to evaluate whether *Rv2983* is necessary for such persistence, the same strains were studied in a model of progressive hypoxia *in vitro*. After 17 days, the change in color of the methylene blue (from blue to yellow) indicated the onset of oxygen deprivation (designated day 0) (Fig. 4C). While there was no difference in the CFU counts between strains at day 0, the viability of the *Rv2983* mutant decreased more rapidly over the ensuing 10 and 21 days (*p* < 0.05 and 0.001, respectively) (Fig. 4C).

Resistance-conferring mutations may confer fitness defects *in vivo*. However, F_420_-deficient mutants have not been well studied in an animal model. In order to understand the effect of *Rv2983* mutations on *M. tuberculosis* virulence *in vivo*, the same strains were subjected to low-dose aerosol infection of BALB/c mice and monitored over the next 4 months. No significant differences in lung CFU counts between the wild-type and the mutant strains were observed (Fig.4D). Similar results were also observed for mouse body, lung and spleen weights (Fig.S5) and lung histopathology, which demonstrated the expected cellular granulomas comprised of histiocytes, foamy macrophages and lymphocytes on day 112 post-infection (Fig. S6). The attenuation of an *Rv2983* mutant in the progressive hypoxia model and the trend towards fewer *Rv2983* mutations in C3HeB/FeJ mice (2/45, 4% of all mutations) compared to BALB/c mice (7/54, 13%) suggests that it may be worthwhile to investigate the role of *Rv2983* in *M. tuberculosis* virulence using C3HeB/FeJ mice in a future study.

### A F_420_-deficient pretomanid-resistant *Rv2983* mutant is hypersusceptible to anti-TB drugs

Previous studies provided evidence that F_420_H_2_ may be necessary for full tolerance to a variety of anti-TB drugs, including isoniazid, rifampin, ethambutol, pyrazinamide, moxifloxacin and clofazimine in *Mycobacterium smegmatis* (33), and isoniazid, moxfloxacin and clofazimine in *M. tuberculosis* (30). In order to confirm that Rv2983 also contributes to tolerance to selected anti-TB drugs in *M. tuberculosis*, we performed time-kill assays exposing the wild-type, the *Rv2983* mutant and the complemented strains to 5-10x MIC concentrations. The mutant proved more sensitive to isoniazid after 4 days of exposure (Fig. 5A). Interestingly, on day 7, the CFU/ml continued to decrease for the mutant, but the wild-type and the complemented strains showed re-growth suggesting that Rv2983 played a role in enabling outgrowth of INH-resistant mutants (Fig. 5A). The *Rv2983* mutant was also hypersusceptible to linezolid, bedaquiline and clofazimine, phenotypes that were fully ameliorated with complementation (Fig. 5B-D).

**Figure 5.**
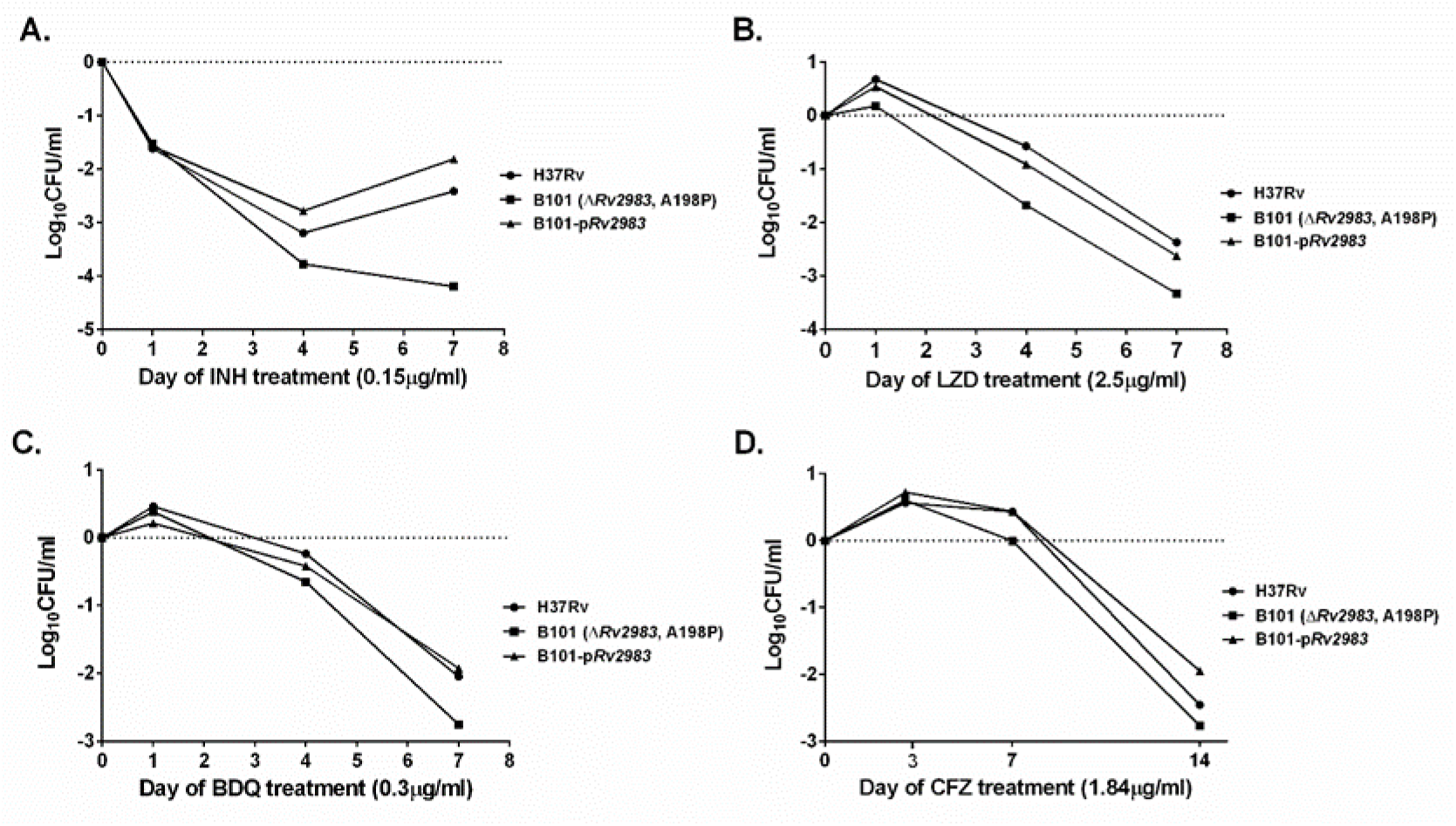
F_420_-deficient pretomanid-resistant *Rv2983* mutant is hypersusceptible to anti-TB drugs. Time-kill kinetics was performed using *Mtb* strains in 7H9 broth containing following drugs: A. INH of 0.15 µg/ml; B. LZD of 2.5 µg/ml; C. BDQ of 0.3 µg/ml; D. CFZ of 1.84 µg/ml. The difference in CFU/ml was calculated based on the CFU/ml at each time point relative to that on day 0 (after subculture of the strains to a drug-containing medium).

### F_420_-deficient pretomanid-resistant mutants are attenuated for growth in the presence of malachite green

Malachite green (MG) is an organic compound used as a selective decontaminant in solid media for culturing *M. tuberculosis*. Previous work using *M. smegmatis* showed that mutations in MSMEG_5126 (homolog of *fbiC*) and MSMEG_2392 (which shares 69% homology with *Rv2983*) reduce the ability to decolorize and detoxify MG, indicating that F_420_ biosynthesis is necessary for this process (21). To evaluate the role of each gene associated with nitroimidazole activation in the resistance to MG, log-phase cultures of selected pretomanid-resistant mutants including the B101 mutant were plated on 7H9 agar supplemented with a range of MG concentrations. All mutants deficient in F_420_ synthesis or F_420_ reduction (*i*.*e*., those with mutations in *fbiA-C, Rv2983* or *fgd*) were more susceptible to MG, while the *ddn* mutant retained the same susceptibility as the wild type H37Rv parent (Fig. 6A). The lability of F_420_H_2_ and lack of a commercial source for F_420_ made it unfeasible to attempt to test whether provision of F_420_H_2_ could rescue the MG-hypersusceptible phenotype of the F_420_H_2_-deficient mutants. However, complementation of *Rv2983* nearly restored the wild-type growth phenotype in the B101 mutant, confirming that *Rv2983* is necessary for the intrinsic resistance of *M. tuberculosis* to MG (Fig. 6B). Interestingly, at MG concentrations above 6 µg/ml, greater recovery was observed when *Rv2983* was expressed behind the native promoter compared to the *hsp60* promoter, suggesting that unknown factors may play a regulatory role in MG detoxification (Fig. 6B). Longer incubation times and plating at higher bacterial density (500 µl rather than 100 µl of cell suspension per plate) significantly increased colony recovery (data not shown).

**Fig. 6.**
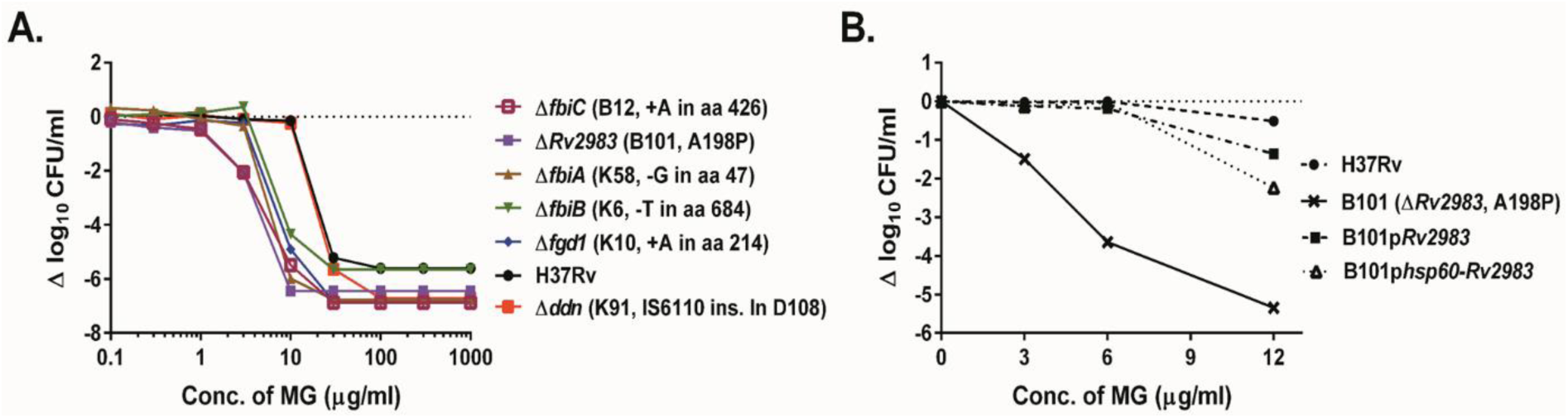
F_420_H_2_-deficient pretomanid-resistant mutants of *M. tuberculosis* are more susceptible to growth inhibition by malachite green. A. Growth of wild-type *M. tuberculosis* on 7H9 agar is inhibited by malachite green (MG) in a concentration-dependent manner. F_420_H_2-_ deficient, pretomanid-resistant *M. tuberculosis* mutants (*fbiA-C, fgd, Rv2983*) are inhibited at lower MG concentrations relative to the wild type and the F_420_H_2_-sufficient, pretomanid-resistant *ddn* mutant. B. Complementation of the B101 mutant with wild-type *Rv2983* restores tolerance to MG after 28 days of incubation.

Because all solid media commonly used to isolate and cultivate *M. tuberculosis* in clinical laboratories contain MG as a selective decontaminant, the increased MG susceptibility conferred by mutations in *fbiA*-*C, Rv2983* and *fgd* could compromise the isolation and propagation (and hence identification) of nitroimidazole-resistant mutants from clinical samples. Commercial 7H10 agar, 7H11 agar and LJ medium contain 0.25, 1 and 400 µg/ml, respectively, of MG. To assess the potential impact of these media on the isolation of an F_420_H_2_-deficient pretomanid-resistant *Rv2983* mutant relative to an F_420_H_2_-sufficient, but still pretomanid-resistant, *ddn* mutant and the pretomanid-susceptible wild type and *Rv2983-*complemented mutant, we inoculated these media in parallel using serial dilutions of each strain. The *Rv2983* mutant exhibited 10 times lower CFU counts relative to other strains after 21 and 28 days of incubation on 7H10 agar plates (*p* <0.01) (Figs. 7A). The result after 35 days of incubation was generally similar between the mutant and the control strains (Fig. 7A). A similar semi-quantitative growth assessment of the *Rv2983* mutant on LJ media compared to other strains including a *ddn* mutant (K91, IS6110 ins in D108) revealed growth inhibition of the *Rv2983* mutant that was ameliorated by increasing the size of the bacterial inoculum from 10^2^ to 10^6^ CFU/ml and increasing the incubation time from 28 to 35 days (Fig. 7C). Interestingly, no difference in growth was found on 7H11 agar (Fig. 7B), despite higher MG concentrations in that medium compared to 7H10.

**Fig. 7.**
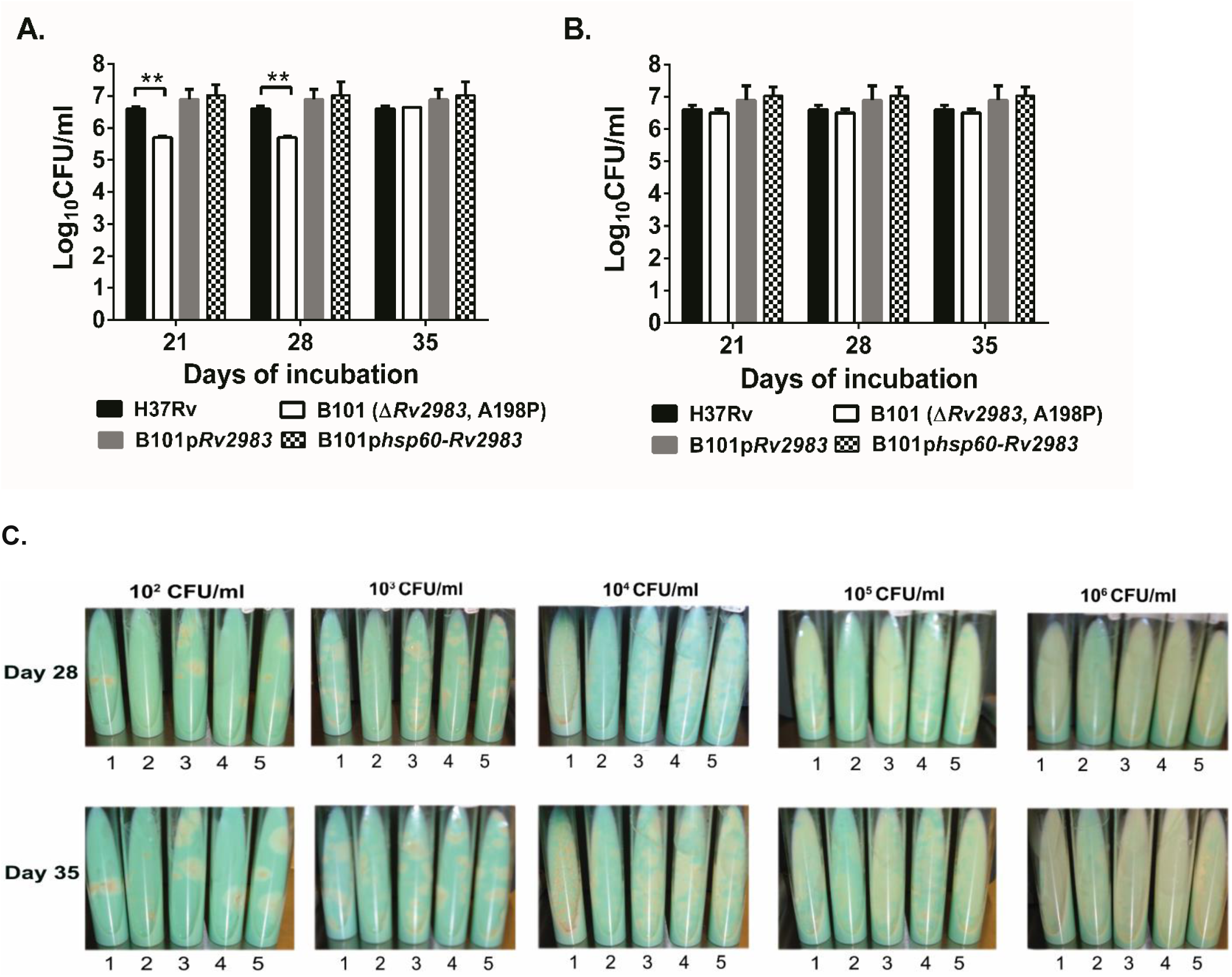
A mutation in *Rv2983* causes growth inhibition on commercial 7H10 agar and LJ slants, but not on commercial 7H11 agar. Aliquots of *M. tuberculosis* cultures were spread on various solid media purchased commercially after serial 10-fold dilutions. A-B. Mean CFU counts on 7H10 (A) and 7H11 (B) agar plates after 21, 28 and 35 days of incubation; C. Colonies on LJ slants inoculated with serially diluted aliquots after 28 and 35 days of incubation. 1: H37Rv wild type; 2: B101 mutant (Δ*Rv2983*, A198P); 3: B101 mutant complemented with *Rv2983* behind the native promoter; 4: B101 mutant complemented with *Rv2983* behind the *hsp60* promoter; 5. K91 mutant (Δ*ddn*, IS6110 ins in D108).

## DISCUSSION

As representatives of one of only two new drug classes approved for use against TB in the last 50 years, delamanid and pretomanid are important and promising new drugs (3, 4, 6, 7, 9, 34) that are increasingly used to treat MDR/XDR-TB. Comprehensive knowledge of the spectrum of mutations conferring resistance to these drugs in *M. tuberculosis in vivo* and the resultant mutant phenotypes is critical for timely and accurate diagnosis of resistance and the design of optimal treatment regimens to promote the safe and effective use of these drugs in clinical settings. The present study reports several important new findings. First, we identified a novel nitroimidazole resistance determinant—loss-of-function mutations in *Rv2983*—that, in the case of our study, explained all of the pretomanid resistance that was not attributable to mutations in the 5 previously described genes. Together these 6 genes comprise a set of non-essential “targets” for spontaneous resistance mutations that is of unprecedented size for a TB drug and results in a relatively lower barrier to resistance compared to most other TB drugs, except perhaps isoniazid. Second, with one exception, *Rv2983* and *fbiB* mutants showed only low-level resistance to delamanid despite high-level resistance to pretomanid. This finding adds to a previous report associating *fbiB* mutations with low-level delamanid resistance (24) and, together with differences in how delamanid resistance has been defined, may explain why neither *fbiB* nor *Rv2983* mutants have yet been associated with delamanid resistance in clinical isolates (29, 35). Third, we provide additional evidence that *Rv2983* is required for F_420_ biosynthesis in *M. tuberculosis* in support of its recently elucidated role as the guanlylyltransferase *fbiD* (20). Finally, we show that *Rv2983* is essential for tolerance of *M. tuberculosis* to MG, a selective decontaminant present in solid media used to cultivate *M. tuberculosis*, and show that clinical microbiology laboratories could encounter difficulties recovering this and other F_420_H_2_-deficient nitroimidazole-resistant mutants from clinical specimens. For reasons that require further exploration, we observed superior recovery of F_420_H_2_-deficient mutants on 7H11 agar compared to 7H10 agar and LJ media, suggesting that 7H11 agar may be the solid medium of choice for identification of nitroimidazole-resistant mutants in clinical and research settings.

Our study provides the first comprehensive analysis of the spectrum of nitroimidazole-resistant mutants selected *in vivo* and, because we used whole genome sequencing, it represents the most comprehensive analysis of pretomanid resistance mutations made to-date. The spontaneous frequency of resistance to nitroimidazoles in *M. tuberculosis* studied *in vitro* has ranged from 1 in 10^5^ to 7 in 10^7^ CFU (22-24, 36-38), which is consistent with our findings in the lungs of untreated BALB/c mice. The large “target size” for mutations in 6 non-essential genes drives this relatively high frequency, which is as high or higher than that for isoniazid and higher than for rifamycins and fluoroquinolones. Our unpublished observations suggest that similar frequencies of nitroimidazole-resistant mutants exist in sputum isolates collected from treatment-naïve, drug-susceptible TB patients. Delamanid-resistant *M. tuberculosis* has been recovered from patients both before and after delamanid treatment (11, 39-41). To date, emergence of resistance has not been described during use of pretomanid in clinical trials, but such use has been restricted to relatively short treatment durations and/or use in combination with highly active companion drugs. Pretomanid resistance has emerged during combination therapy in mouse models (3, 42). Thus, the relatively high frequency of spontaneous mutations conferring nitroimidazole resistance and available pre-clinical and clinical data underscore the importance of making validated DST for this class widely available as clinical usage expands. Moreover, our finding also emphasizes the importance of using nitroimidazoles in regimens with other effective anti-TB drugs to which infecting strains are susceptible, ideally taking advantage of the hypersusceptibility of F_420_H_2_-deficient mutants to many anti-TB drugs, as shown here and elsewhere to restrict their selective amplification. Indeed, the use of pretomanid in highly active regimens under clinical trial conditions may be an important reason for the absence of treatment-emergent resistance to date.

The lungs of TB patients feature a heterogeneous array of lesion types, which possess diverse immune responses and cause differences in drug penetration (25, 26). C3HeB/FeJ mice develop caseating lung lesions and BALB/c mice form largely cellular lesions in response to *M. tuberculosis* infection. We observed selective amplification of F_420_H_2_-deficient mutants in mice over a range of pretomanid doses that included doses producing much higher drug exposures than those produced in patients. Amplification was especially pronounced at higher drug doses, which eliminated the nitroimidazole-susceptible population more rapidly, and in C3HeB/FeJ mice. Our finding suggests that microenvironments in C3HeB/FeJ mice favor the selective amplification of nitroimidazole-resistant mutants. Further study is needed to explain this finding, but pretomanid is expected to penetrate well into necrotic lesions and to exert activity under relatively hypoxic conditions. The more rapid selection of resistance in C3HeB/FeJ mice has been observed for other drugs and may have more to do with the larger bacterial loads and reduced host immune pressure in the caseating lesions. Fortunately, our study and others show that F_420_ is crucial for mycobacterial tolerance to a range of antimicrobial compounds and that F_420_H_2_-deficient mycobacterial strains are more susceptible to first-line and second-line anti-TB drugs such as isoniazid, rifampin, pyrazinamide, ethambutol, moxifloxacin, bedaquiline, linezolid, clofazimine and other compounds including MG (30, 33). Combining pretomanid and delamanid with these drugs can be expected to counter the selection of F_420_H_2_-deficient nitroimidazole-resistant sub-populations by killing them more rapidly than wild type *M. tuberculosis* sub-populations, as suggested by recent preclinical studies (3).

Previous work identified 5 genes (*fbiA-C, fgd*, and *ddn*) involved in the activation pathway of nitroimidazole prodrugs in which mutations may confer drug resistance in *M. tuberculosis* complex (17, 19, 22-24, 38). Like the *in vitro* study by Haver *et al* (23), we found that isolated mutations in *fbiA-C, fgd*, or *ddn* explained the majority of the pretomanid-resistant isolates we selected. However, whereas their study left 17% of resistant isolates unexplained, we found that all of the remaining resistant isolates in our study, representing 9% of the total number of unique mutations, harbored mutations in *Rv2983*, a gene not previously implicated in nitroimidazole resistance. Indeed, the proportion of resistant isolates explained by *Rv2983* (9%) was similar to the proportion explained by *fbiA* (15%) and *ddn* (12%) mutations, which lagged only mutations in *fbiC* (56%) as the predominant cause of pretomanid resistance in our mice. Thus, the identification of *Rv2983* mutations should be included in rapid molecular DSTs and algorithms for the diagnosis of nitroimidazole resistance from genome sequence data. The 10 mutations in *Rv2983* identified in this study (Table S2-4) represent the first step in the process of identifying specific resistance-conferring mutations to inform test development. Although the *Rv2983* mutants caused a smaller upward shift in the delamanid MIC compared to the pretomanid MIC, our complementation study proves that *Rv2983* is also required for efficient delamanid activation. The delamanid MIC of 0.064 µg/ml against the mutant was still higher than the recently proposed critical concentration of 0.016 µg/ml (29). Interestingly, all of our *fbiB* mutants also demonstrated only low-level resistance to delamanid (2-8x increase in MIC) despite high-level pretomanid resistance (32-128x increase in MIC. Such low-level delamanid resistance with mutation of *fbiB* was also observed in *M. bovis* BCG by Fujiwara et al (24). This finding suggests that delamanid may be less likely to select such mutants in these genes and may retain more activity than pretomanid against these mutants due to the smaller selection window between the mutant and wild-type MICs. The reason for the differential impact of these mutations on pretomanid and delamanid susceptibility remains unexplained. However, it is conceivable that *M. tuberculosis* can utilize even relatively modest levels of Fo, F_420_-0 or dehydro-F_420_-0 produced in the absence of FbiD or FbiB to activate delamanid. Apparently, this is not the case for pretomanid, which may be due in part to differences in the chemical structure of the two drugs and the impact on the efficiency of Ddn-mediated activation. This warrants further investigation.

Our WGS results confirm and significantly extend prior *in vitro* work demonstrating the remarkable diversity of mutations capable of conferring high-level nitroimidazole resistance. Among the 99 unique mutations we identified in 47 mice, only 3 mutations (K9N in *fgd*, R322L in *fbiC* and Q120P in *fbiA*) were found in more than one mouse and each mouse generally hosts 1 to 4 unique mutations. Furthermore, by comparing the 99 unique mutations observed in our mice with the 151 unique mutations selected *in vitro* (23), the same mutation occurred only twice. Thus, each of the 6 genes now implicated in nitroimidazole resistance appears to be devoid of “hot spots” for such mutations. The unprecedented number and diversity of resistance-conferring mutations demonstrated for nitroimidazole drugs here and by Haver *et al* (23), clearly challenges the development and interpretation of rapid molecular susceptibility tests, especially considering that polymorphisms in nitroimidazole resistance genes that represent phylogenetic markers but do not confer pretomanid resistance are well-described (43, 44). A similar situation exists for *pncA* mutations and pyrazinamide (PZA) resistance, where an efficient, yet comprehensive method based on saturating mutagenesis for distinguishing single nucleotide polymorphisms conferring resistance was recently described (45). A similar analysis of substitutions in the 6 genes related to nitroimidazole resistance would similarly advance the development of DST using genome sequencing technology.

Bashiri et al recently revised the F_420_ biosynthetic pathway based on biochemical evidence that Rv2983 catalyzes production of the guanylated PEP moiety that is used with Fo by FbiA to produce dehydro-F_420_-0 (20). Using overexpression of *Rv2983* in *M. smegmatis* and *M. tuberculosis Rv2983* mutants, we show that expression of *Rv2983* is necessary for efficient conversion of Fo to F_420_, and thereby providing the first evidence that *Rv2983* is necessary for this step in the pathogen *M. tuberculosis* and adding to previous evidence that its ortholog MSMEG_2392 is involved in F_420_ biosynthesis in *M. smegmatis* (21). The validity of the method used in this study for detection of F_420_ and Fo was demonstrated by showing the expected results with two pretomanid-resistant strains, KA016 and KA026, harboring mutations in *fbiC* and *ddn*, respectively.

Our findings regarding the heightened susceptibility of F_420_H_2_-deficient mutants to MG pose a previously unappreciated challenge to the development and use of phenotypic testing methods. Indeed, we observed reduced or delayed recovery of a nitroimidazole-resistant *Rv2983* mutant on commercial 7H10 and LJ media that include MG as a selective decontaminant (46). Since *fbiA-C* and *fgd* mutants exhibited similar hypersusceptibility to MG in 7H9 agar supplemented with MG, their recovery on 7H10 and LJ is also likely to be affected. Although liquid culture media such as MGIT media are increasingly used in clinical microbiology laboratories, the selective growth inhibition of nitroimidazole-resistant strains on solid media that are still commonly used in clinical microbiology laboratories around the world for isolation and subculture of *M. tuberculosis* raises serious concern that their recovery from clinical specimens such as sputum, could be impaired, especially for isolates comprised of mixed wild-type and resistant populations. This concern is further amplified by the common practice of performing susceptibility testing (including molecular testing), not on primary samples but, on isolates that have been sub-cultured one or more times on solid media. Such practices may drastically reduce the proportion of (or eradicate) F_420_H_2_-deficient mutants present in the original sample. In addition, efforts to develop MG decolorization assays for detection of drug-resistant TB are expected to be fruitless for these mutants (47-50). We did not determine the basis for the greater recovery of F_420_H_2_-deficient mutants on 7H11 vs. 7H10 media despite 4x higher total MG concentrations in the former. The principal differences between these media are the presence of pancreatic digest of casein in 7H11 and lower concentrations of magnesium sulfate countered by the addition of copper sulfate, zinc sulfate and calcium chloride in 7H10. Although this issue clearly requires further study, we presently believe that 7H10 and LJ should not be employed for phenotypic nitroimidazole susceptibility testing and that primary isolation or subculture of any isolate on such media prior to either phenotypic or genotypic susceptibility testing should be avoided whenever possible. When it cannot be avoided, larger inoculum sizes and longer incubation times may increase recovery on 7H10 and LJ. Based on our study, 7H11 agar appears to be the preferred solid medium for recovery of F_420_H_2_-deficient nitroimidazole-resistant *M. tuberculosis*.

In conclusion, using BALB/c and C3HeB/FeJ mice and WGS, we characterized the pretomanid dose-response relationships for bactericidal effect and suppression of drug-resistant mutants and profiled the genetic spectrum of pretomanid resistance emerging *in vivo*. A novel resistance determinant, Rv2983, was identified as essential for F_420_ biosynthesis and activation of the novel pro-drugs delamanid and pretomanid. Furthermore, we provide evidence that F_420_H_2_-deficient, nitroimidazole-resistant *M. tuberculosis* mutants are hypersensitive to MG, raising concern that using MG-containing medium could compromise the isolation and propagation of *M. tuberculosis* from clinical samples and therefore hinder the clinical diagnosis of nitroimidazole resistance. These findings have important implications for both genotypic and phenotypic susceptibility testing and treatment strategy to detect and eliminate nitroimidazole resistance, which will be of increasing importance as wider use of delamanid and pretomanid ensues. More comprehensive understanding of the spectrum of resistance mutations that emerge during treatment with new drugs *in vivo* should be considered as an integral part of TB drug development prior to clinical application.

## MATERIALS AND METHODS

### Bacterial strains, media, antimicrobials and reagents

Wild type *M. tuberculosis* H37Rv (ATCC 27294) was mouse-passaged, frozen in aliquots and used in all the experiments. The wild type *M. smegmatis* strain mc^2^ 155 was obtained from the stock in the lab. Unless stated otherwise, Middlebrook 7H9 medium (Difco, BD) supplemented with 10% oleic acid-albumin-dextrose-catalase (OADC) complex (BD), 0.5% glycerol and 0.05% Tween 80 (Sigma-Aldrich) (7H9 broth) was used for cultivation. Dubos Tween Albumin Broth (BD Difco) supplemented with the hypoxia indicator methylene blue (Sigma-Aldrich, 500 mg/L) was prepared for the progressive hypoxia study. Middlebrook 7H10 agar and selective 7H11 agar (Difco, BD), prepared from powder and containing 10% OADC and 0.5% glycerol, were used for comparison of strain recovery on commercially available agar plates. Lowenstein Jensen (LJ) slants were purchased from BD. Pretomanid, delamanid and bedaquiline were kindly provided by the Global Alliance for TB Drug Development (New York, NY). Isoniazid, linezolid, clofazimine and menadione were purchased from Sigma-Aldrich.

## Mouse infection models and pretomanid treatment

All animal procedures were approved by the Animal Care and Use Committee of Johns Hopkins University. Aerosol infections were performed using the Inhalation Exposure System (Glas-col Inc., Terre Haute, IN), as previously described (51). Briefly, 6-week-old female BALB/c mice (Charles River, Wilmington, MA) and C3HeB/FeJ mice (Jackson Laboratories Bar Harbor, ME) were infected with a log phase culture of *M. tuberculosis* that was grown in 7H9 broth to O.D.600_nm_ = 1.0 and then diluted in the same medium prior to infection to deliver 50-100 CFU to the lungs. Pretomanid was formulated for oral administration as previously described (37). Beginning 8 weeks after aerosol infection, mice were randomly allocated into groups and treated once daily (5 days per week) for up to 8 weeks with pretomanid at doses of 10, 30, 100, 300 and 1000 mg/kg. Untreated mice were sacrificed on the day after aerosol infection and on the day of treatment initiation to determine the number of CFU implanted in the lungs and pretreatment CFU counts, respectively. Additional mice were sacrificed after 3 and 8 weeks of treatment to evaluate the treatment response. Serial 10-fold dilutions of lung homogenates were plated on 7H11 agar. Week 8 samples including those from untreated mice were also plated in parallel on 7H11 plates containing 0.25, 1 and 10 µg/ml of pretomanid to quantify the resistant CFU. Plates were incubated at 37°C for 28 days before final CFU counts were determined.

### Whole genome sequencing

For each mouse lung that yielded growth on pretomanid-containing plates, individual colonies and, for a subset of mice, pools of up to 15 colonies, were randomly selected from pretomanid-containing plates and sub-cultured in 7H9 broth prior to extraction of genomic DNA using the cetyltrimethylammonium bromide (CTAB) protocol (52) and vortexing (Genegate, Inc.). 2-3 μg of genomic DNA was sheared by a nebulizer to generate DNA fragments. The DNA library was prepared using a genomic DNA sample preparation kit (Illumina, Inc.), in which adapter-ligated DNA fragments were 250-350 bp in length, and carried out on an Illumina HiSeq 2500 (Illumina, Inc). The sequencer was operated in paired-end mode to collect pairs of reads of 72-bp from opposite ends of each fragment. Image analysis and base-calling were done by using the Illumina GA Pipeline software (v0.3). The reads that were generated for each strain were aligned to the reference genome of *M. tuberculosis* H37Rv (53). Based on alignment to the corresponding region in the reference genome, single nucleotide polymorphism (SNP), insertion and deletion were identified on the genome of resistant strains by using a contig-building algorithm to construct a local ∼200 bp sequence spanning the site of mutagenesis (54). Distribution of mutation type and mutation frequency in genes involved in nitroimidazole resistance was calculated by counting the total number of unique mutations isolated from each mouse in the same treatment group.

### Complementation of an *Rv2983* mutation

A 1,044-bp DNA fragment containing the open reading frame (ORF) of the wild type *Rv2983* gene, including 340 bp of 5’-flanking sequence and 59 bp of 3’-flanking sequence, was PCR-amplified from *M. tuberculosis* H37Rv genomic DNA using primers *Rv2983*-1F and *Rv2983*-1R (Table S1). The *Rv2983* PCR product was ligated into XbaI-digested *E. coli-*mycobacterium shuttle vector pMH94 (28) using NE builder HiFi DNA assembly kit (NE Biolabs) to generate the recombinant pMH94-*Rv2983* vector. Similarly, a 388-bp DNA fragment containing the *hsp60* promoter and a 645-bp DNA fragment of *Rv2983* open reading frame were amplified from *M. tuberculosis* H37Rv genomic DNA using primer sets *hsp60*-F and *hsp60*-R and *Rv2983*-2F and *Rv2983*-2R, respectively (Table S1), and ligated into XbaI-digested *E. coli-*mycobacterium shuttle vector pMH94 to yield pMH94-*hsp60*-*Rv2983*. A small amount of ligation reaction was transferred into *E. coli* competent cells, followed by DNA sequencing of the inserts in the corresponding recombinants. The recombinants pMH94-*Rv2983* and pMH94-*hsp60*-*Rv2983* were electroporated into competent cells of *Rv2983* mutant strain BA_101 (B101), harboring an A198P substitution, to enable selection of complemented candidates B101p*Rv2983* and B101p*hsp60-Rv2983* on 7H10 agar containing 25 µg/ml of kanamycin. To confirm the complementation genetically, Southern blotting was performed using a digoxigenin (DIG) DNA labeling and detection kit according to the manufacturer’s protocol (Sigma). Briefly, a 448-bp *Rv2983* probe was generated by addition of DIG-dUTP (Sigma) to PCR reactions containing primer pairs *Rv2983*-3F and *Rv2983*-3R (Table S1). Acc65I-digested (NE biolabs) genomic DNA of the wild type, the B101 mutant and the B101p*Rv2983* and B101p*hsp60-Rv2983* complemented strains was separated on agarose gel and transferred onto positively-charged nylon-membrane (GE). After pre-hybridization, the membrane was hybridized with the DIG-labeled *Rv2983* probe at 68°C overnight, followed by addition of anti-DIG alkaline phosphatase conjugate. After stringent washes, the membrane was incubated with the chemiluminescence substrate disodium 3-(4-methoxyspiro {1,2-dioxetane-3,2(5′-chloro)tricycloecan}-4-yl)phenyl phosphate (CSPD) and exposed on X-ray film in a dark room prior to development using a developer (AFP imaging)(27).

### MIC determination

MICs were determined using a broth macrodilution assay. Log-phase cultures were adjusted to achieve a bacterial density of approximately 10^5^ CFU/ml when added to conical tubes containing complete 7H9 broth without Tween 80 and with or without either pretomanid or delamanid in concentrations ranging from 0.015 to 32 µg/ml or from 0.001 to 16 µg/ml, respectively. Drugs were initially dissolved in dimethylsulfoxide (DMSO) (Sigma) prior to further dilution in 7H9 broth. Cultures were incubated at 37°C for 14 days. MIC was defined as the lowest drug concentration that inhibited visible *M. tuberculosis* growth (55, 56). The experiments were performed at least twice for each strain.

### Time-kill assays

Mid-log-phase cultures of *M. tuberculosis* were diluted to OD_600nm_ of 0.001 (about 10^5^ CFU/ml) in 3 ml of 7H9 broth, exposed to isoniazid, linezolid, bedaquiline or clofazimine, and then incubated in a 37°C shaker for 7 or 14 days. Aliquots were plated on 7H11 agar after serial dilutions and incubated for 21 days at 37°C prior to CFU counting. The experiments were performed twice.

### Oxidative stress and progressive hypoxia assays

To observe the response to menadione-induced oxidative stress, mid-log phase cultures of *M. tuberculosis* were diluted to OD_600nm_ of 0.01 (about 10^6^ CFU/ml) in 3 ml of 7H9 broth containing varying concentrations of menadione or no menadione. The cultures were incubated in a 37°C shaker for 6 days. To study survival under progressive hypoxia, the mid-log phase cultures were diluted to OD_600nm_ of 0.001 (about 10^5^ CFU/ml) in 20 ml of Dubos Tween Albumin Broth with methylene blue (500 mg/L) in rubber-cap test tubes (25mmX125mm) with sterile magnetic stir bars. The tubes were sealed and incubated upright on a magnetic platform at 37°C until the methylene blue dye changed to yellow, indicating the depletion of oxygen, and then incubated for an additional 21 days after the color change (57). Samples from various time points were collected from the above cultures and plated on 7H11 agar plates after serial dilutions followed by 21 days of incubation at 37°C before CFU counting. In the progressive hypoxia assay, samples were taken by carefully inserting a syringe needle through the rubber stopper to avoid introducing oxygen to the cultures.

### Virulence assessment in BALB/c mice

Female BALB/c mice (6-8 weeks of age) (Charles River Labs) were aerosol-infected with approximately 100 CFU of *M. tuberculosis* using the Inhalation Exposure System (Glas-Col). After infection, groups of 4–5 mice were sacrificed on day 1 and at designated time points thereafter. Lungs and spleens were removed aseptically. The weights of body, lung and spleen were measured and recorded. The upper lobe of the left lung was removed, fixed in paraformaldehyde and processed for histological examination by hematoxylin and eosin staining. After serial dilution the homogenates of the remaining lung tissues were plated on Middlebrook 7H11 selective agar plates (Thermo Fisher Scientific) and incubated at 37°C for 28 days prior to CFU counting.

### Construction of recombinants overexpressing *Rv2983*, with or without *fbiC*, in *M. smegmatis*

A 645-bp DNA fragment containing the *Rv2983* ORF was PCR-amplified from *M. tuberculosis* H37Rv genomic DNA using primers *Rv2983*-4F and *Rv2983*-4R (Table S1). The amplified PCR product was ligated into the NdeI- and PacI-digested *E. coli-*mycobacterium shuttle vector pYUBDuet (58) using NE builder HiFi DNA assembly kit (NE Biolabs) and then transferred into Turbo-competent *E. coli* cells (NE Biolabs) prior to plating on LB agar plates containing 100 µg/ml of hygromycin B for selection of recombinants. The *Rv2983* PCR product was also similarly ligated into the same NdeI- and PacI-digested pYUBDuet vector harboring *fbiC* (termed p*fbiC*) (58) to overexpress both *Rv2983* and *fbiC*. After confirmation by restriction digestion and DNA sequencing, the constructs were electroporated into competent *M. smegmatis* cells prior to selecting recombinants on 7H10 agar plates containing 100 µg/ml of hygromycin B. PCR amplification of the hygromycin resistance gene with primers hyg-F and hyg-R (Table S1) was used to confirm the inserts on the *M. smegmatis* genome. pYUBDuet and pYUBDuet harboring *fbiA, fbiB and fbiC* (termed p*fbiABC*) (58) were also transferred into competent *M. smegmatis* cells to serve as controls.

### Measurement of Fo and F_420_

Extraction of Fo and F_420_ was performed in *M. smegmatis* and *M. tuberculosis* strains according to a previous study (58), with minor modifications. Briefly, *M. smegmatis* strains harboring different constructs and pYUBDuet were grown in 7H9 broth in a shaker to mid-log phase (O.D._600nm_ = 0.7-1.0), followed by induction using 1mM isopropyl β-D-1-thiogalactopyranoside (IPTG) for 6 and 26 hours. After centrifugation for 15 min at 16000 x g, the supernatants were removed for detection of Fo, which is principally found in culture supernatant whereas F_420_ with 5 or 6 glutamate residues is largely retained inside cells (16, 58, 59). The cell pellets were washed with 25mM sodium phosphate buffer (pH 7.0) and re-suspended at 100 mg/mL in the same buffer, then autoclaved at 121°C for 15 min. After centrifugation at 16000 x g for 15 min at 4°C, the cell extracts were harvested for detection of F_420_ (58). Fluorescence of the supernatant and cell extracts was measured using an excitation wavelength of 410 nm and an emission wavelength of 465 nm. Fluorescent signals of Fo were normalized using the O.D. at 600nm. The small portion of Fo (1-7%) retained inside cells was ignored when quantifying F_420_ in cell extracts (60). Relative fluorescent signals were calculated in *M. smegmatis* harboring each of recombinants relative to pYUBDuet alone. Similarly, cell extracts and supernatant were also extracted from *M. tuberculosis* strains grown in 7H9 broth for 6 days at initial O.D._600nm_ of 0.1. Relative fluorescent signals of F_420_ and Fo were calculated using cell extracts and supernatant relative to 25 mM phosphate buffer and 7H9 broth, respectively. *M. smegmatis* harboring pYUBDuet-*fbiABC* was used as a positive signal control for Fo and F_420_ due to their commercial unavailability (58). The experiment was repeated twice.

### Malachite green susceptibility testing

7H9 media supplemented with 10% OADC, 0.5% glycerol, 1.5% Bacto™ Agar (BD) and malachite green (MG) oxalate (Alfa Aesar) was used to prepare solid 7H9 media with differing MG concentrations. *M. tuberculosis* strains were grown to mid-log phase and diluted to OD_600nm_ = 0.1 in 7H9 broth before serial 10-fold dilutions were plated in 100 or 500 µl aliquots on 7H9 agar containing MG concentrations of 0, 0.1, 0.3, 1, 3, 10, 30, 100, 300, 1000 µg/ml or 0, 3, 6, 12 µg/ml. CFU were counted after 28, 35 and 49 days of incubation. The same cultures were also plated on 7H10 and 7H11 agar plates and LJ slants. Serially diluted cultures were inoculated onto LJ slants using calibrated disposable inoculating loops (10 µl per loop, BD) as one loop per LJ slant. Plates were incubated at 37°C for 21, 28 and 35 days for CFU counts. Colony size was observed weekly until day 35, beginning 21 days after plating. The experiment was repeated two times under similar conditions.

## Statistical analysis

Log_10_-transformed CFU counts, fold-change values of gene expression and absorbance (A_410_) values of fluorescent signals were used to calculate means and standard deviations for each data set. Differences between means were compared by the Student’s *t* test in Microsoft Excel. Differences in mutation frequencies between two mouse models were evaluated by Fisher’s exact test in GraphPad Prism 6. A *p*-value of < 0.05 was considered statistically significant.

## Acknowledgements

The Global Alliance for TB Drug Development kindly provided pretomanid and delamanid under a material transfer agreement. The authors gratefully acknowledge Anna Upton and Juliano Timm for critical reading of the manuscript.

## Author contributions

D.R. and E.L.N. conceived the study and designed the experiments. S.L. and J-P.L. assisted with the design and conduct of the *in vivo* experiment. Whole genome sequencing was performed and analyzed by T.I. and J.S., and the results were further analyzed by D.R. and E.L.N. *In vitro* experiments were performed and analyzed by D.R., K.S., J.L. and E.L.N. The manuscript was drafted by D.R. and E.L.N. with critical input from T.I., J.S., and G.B.

## Data and materials availability

All data necessary for evaluation of the conclusions are present in the paper and/or the Supplementary Materials.

## SUPPLEMENTARY MATERIALS

**Table S1**. List of the primers used in the study

**Table S2**. WGS results of 82 individual pretomanid-resistant colonies from BALB/c mice

**Table S3**. WGS results of 54 individual pretomanid-resistant colonies from C3HeB/FeJ mice

**Table S4**. WGS results of 25 pooled pretomanid-resistant isolates selected from BALB/c and C3HeB/FeJ mice

**Table S5**. Distribution of mutation types and frequencies in the genes associated with pretomanid resistance BALB/c and C3HeB/FeJ mice

**Figure S1**. Complementation of B101 mutant with *Rv2983*. A. Schematic diagram of genomic DNA of *M. tuberculosis* strains after digestion with restriction enzyme Acc65I; B. Result of southern blot confirmed expected DNA fragments after Acc65I digestion using DIG-labeled *Rv2983* probe (H37Rv: 6.3 kb; Rv2983 mutant: 6.3 kb; complemented strains: 6.3 and 3.5 kb).

**Figure S2**. Expression of *Rv2983* and other genes involved in nitroimidazole activation. A. Expression of *Rv2983* and other genes involved in nitroimidazole activation is higher in the *Rv2983* mutant B101 relative to the wild-type H37Rv after 4 days of incubation in 7H9 broth; B. *fbiC* expression is dramatically lower in the *fbiC* mutant KA026 relative to the wild-type after 2 days of incubation in 7H9 broth; C. A faint band representing the 937-bp *fbiC* DNA fragment is evident in the sample from the KA026 mutant (lane 2) relative to that in H37Rv (lane 3). Lane 1 is the 1-kb DNA marker.

**Figure S3**. Complementation of the B101 mutant with wild-type *Rv2983* restores tolerance to MG. The proportional recovery of the mutant on 6 µg/ml of MG increases with the volume of culture plated and the duration of incubation: 28-day incubation of 500 µl (A) aliquots/plate; 35-day incubation of 500 µl (B) aliquots/plate.

**Figure S4**. The growth kinetics of the wild-type H37Rv, the Rv2983 mutant B101 and the complemented strains (B101-p*Rv2983* and B101-p*hsp60-RV2983*) in 7H9 broth.

**Figure S5**. Gross examination of aerosol-infected BALB/c mice sacrificed at different time points. A. Body weight; B. Lung weight; C. Spleen weight.

**Figure S6**. Histopathological examination of lung tissues on day 112 post-infection (hematoxylin & eosin staining; 200X magnification).

